# A grid-like basis for affective space in human ventromedial prefrontal cortex

**DOI:** 10.64898/2026.04.30.721013

**Authors:** Yumeng Ma, Philip A. Kragel

## Abstract

Human emotional experiences vary widely in everyday life, yet they are systematically organized within a continuous affective space defined by valence and arousal. Although this circumplex organization has shaped decades of research, its neural implementation remains unclear. Here, we asked whether abstract emotion concepts are encoded using grid-like coordinate principles in the human brain. During fMRI, participants tracked trajectories through affective space conveyed through morphs of facial expressions and sequences of phrases. We found that ventromedial prefrontal cortex (vmPFC) encoded affective space with robust hexadirectional modulation characteristic of grid-like coding. This coordinate representation generalized across stimulus modalities, indicating abstraction beyond perceptual features. The grid-like signal reflected a stable metric organization of emotion concepts independent of overt behavior, whereas distance-related signals in vmPFC predicted trial-by-trial affective judgements. Together, these findings suggest that emotion knowledge is structured by a grid-like coordinate system in the human brain that can support predictive inference and flexible decision-making.

## Introduction

Humans navigate a wide range of events in everyday life, from pleasant, calm conversations to tense, conflict-laden encounters. These episodes are complex, varying in the external situations, internal bodily sensations, and potential actions that define them. To behave adaptively in such environments, the brain is thought to organize accumulated knowledge into structured representations that support prediction, inference, and flexible behavior [1,2]. Converging evidence suggests that diverse forms of knowledge, including visual objects [3], people [4], odors [5], choice options [6], and action plans [7], are encoded in a map-like way in medial temporal and prefrontal systems. Such representations embed specific experiences within continuous spaces, allowing relationships among states to be expressed geometrically, enabling generalization beyond immediately available sensory inputs [2].

Among the many domains of knowledge that guide flexible behavior, information about emotional events is particularly important for survival and well-being [8]. Decades of behavioral research indicate that individuals organize emotion knowledge in a two-dimensional affective space defined by valence and arousal [9,10]. This space is consistently observed across judgments of facial [11], vocal expressions [12], emotion words [9,13], and self-reported feelings [14], suggesting that individuals abstract meaning from diverse sensory experiences to represent a shared structure of affect. This framework, known as the affective circumplex, has been proposed to reflect the cognitive structure laypeople use to organize knowledge about emotions and guide behavior [9].

Despite the robustness of the affective circumplex structure in behavioral data, its neural basis remains unclear. Contemporary theories explaining the brain basis of emotion have proposed that the default network constructs predictive models of the world to anticipate the needs of an organism in context [15,16]. Within this network, ventromedial prefrontal cortex (vmPFC) has been implicated in integrating long-term memory with information about the self and current goals to generate value-based predictions that guide decision-making [17,18]. These accounts explain how affective meaning can be constructed flexibly across situations, yet they do not specify the representational format used to organize emotional knowledge. As a result, they do not explain why circumplex structure emerges so consistently across modalities, tasks, and individuals [10].

One possibility is that hippocampal-prefrontal components of the default network represent emotion knowledge in a map-like way [19]. In this account, neural activity in vmPFC reflects the relational structure among emotional events, enabling flexible generalization across contexts. However, relational structure alone does not specify the computational format of a map [20]. Thus, it remains unclear whether affective space is implemented in a coordinate-based metric system. In spatial and conceptual domains, such a metric structure is supported by grid-like codes: when participants navigate physical and abstract spaces, fMRI signals in vmPFC and entorhinal cortex exhibit hexadirectional modulation—a six-fold periodic pattern aligned to movement in space [21]. Similar grid-like coding has been observed in vmPFC during economic decision-making involving an abstract value space [22]. Moreover, vmPFC activity in response to emotional films can be predicted based on regularities in emotion transitions consistent with structural generalization [23]. However, direct evidence that affective space itself is represented using grid-like codes remains lacking.

To test whether affective space is implemented using a grid-like basis, here we conducted an fMRI experiment in which participants (*n* = 18) viewed emotion-laden stimuli and judged how they varied along dimensions of valence and arousal. During scanning, participants were presented with morphs of prototypical facial expressions and sequences of phrases (Fig. 1a). Each trial consisted of a single trajectory defined by start- and end-points in affective space. We created stimuli to provide broad coverage across the affective space and validated them via post-scan ratings of valence, arousal, and emotion categories (Fig. 1b). On each trial during scanning, participants indicated whether the valence or arousal of the stimulus increased, decreased, or stayed the same. We varied the probed dimension randomly, requiring participants to track changes in both valence and arousal throughout the task.

**Figure 1.**
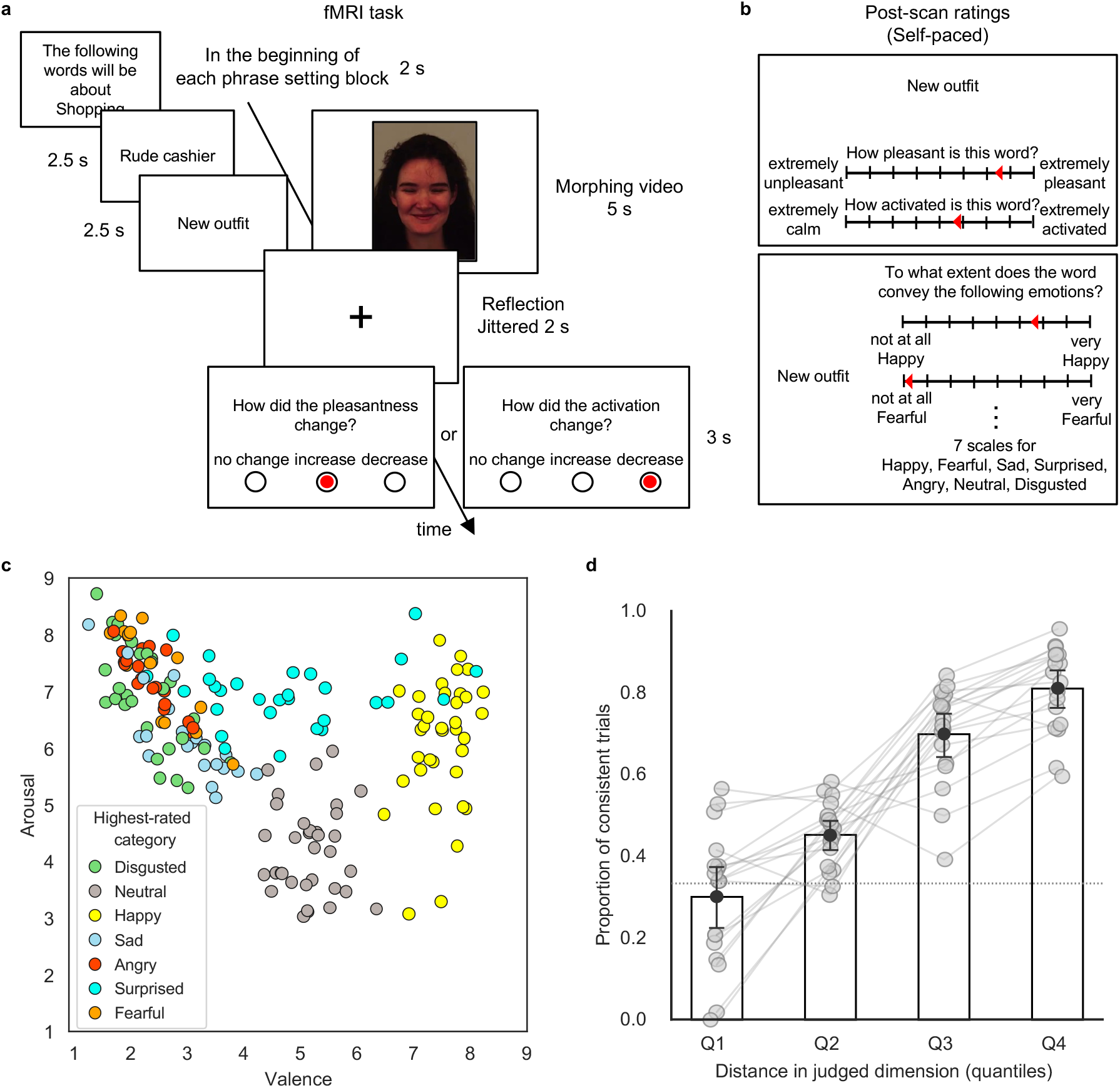
Experimental paradigm and behavior. **a**, Schematic of a single trial in the two versions of the task (phrase and face). Each trial began with a sequence of two phrases from the same setting, or a morph between two facial expressions of the same identity. After a jittered fixation period, participants had 3 s to indicate whether valence or arousal had increased, decreased, or stayed the same. The probed dimension was split evenly across trials and revealed only on the response screen, so participants could not anticipate which dimension would be queried while viewing the stimuli. The face stimulus shown here is from the Karolinska Directed Emotional Faces database [50] (image ID: F04HAS). **b**, Rating screens for one example phrase. After scanning, participants rated each phrase and facial expression on valence and arousal, followed by ratings of seven emotion categories, using continuous scales. **c**, Group-average valence-arousal ratings. Each point represents one stimulus and is colored according to the emotion category with the highest rating. **d**, Proportion of trials in which in-scanner judgments were consistent with the group-average post-scan ratings across four quantiles of absolute distance in the judged dimension. The dotted line indicates chance agreement. Points represent individual participants (*n* = 18 independent participants), and points from the same participant are connected by gray lines. Bars show group means with 95% confidence intervals.

By combining two perceptually distinct stimulus modalities with diverse affective trajectories, this design allowed us to test whether abstract emotion concepts are organized using a modality-invariant coordinate system. If this is the case, we would expect two outcomes. First, post-scan ratings should recover the geometric organization of affect that defines the circumplex. Second, BOLD signals in vmPFC should exhibit hexadirectional modulation as a function of trajectory direction through that space, reflecting grid-like coding that generalizes across stimulus modalities and is independent of task demands.

### Grid-like coding of affective space in vmPFC

A map-like organization of affect entails that judgments of emotional stimuli are based on a Euclidean space in which emotion concepts are embedded [9]. Indeed, analyses on the post-scan ratings revealed a two-dimensional affective structure broadly similar across participants (S1-S3 Fig and S1-S2 Table), stimulus modalities (S4 Fig), and rating types (Fig. 1c and S5 Fig). Although each of these factors could introduce meaningful variability, the dominant structure in our data resembled a circumplex, with positive categories (e.g., happy, surprise) clustering toward high valence, negative categories (e.g., fear, sadness, anger, disgust) toward low valence regions, and arousal increasing from neutral to highly activating categories like surprise and fear.

To examine whether this structure aligned with choice behavior during the fMRI scan, we assessed the consistency of in-scanner judgments of valence and arousal change with the affective structure derived from post-scan ratings. We found that as the distance between stimuli in the judged dimension increased, so did the agreement between individual choices during scanning and the group-average post-scan ratings (logistic regression: *β* = 0.9137, *SE* = 0.1212, *z* = 7.540, 95% CI [0.6770, 1.1599], *p* < .0001; overall proportion of consistent choices: range = 50.96-66.35%; median = 56.97%, IQR = 7.21%, chance = 33.33%; Fig. 1d). These behavioral results indicate that participants made judgements consistent with a common, circumplex organization of emotion knowledge during scanning, enabling tests of grid-like structure in fMRI measures.

When participants viewed face morphs and phrase sequences, transitions between emotion concepts could be represented as movement through affective space. If the basis for navigating this space is implemented in grid-like population codes, trajectories aligned with a common grid orientation should evoke similar fMRI patterns separated by 60° rotational symmetry. To test this prediction, we estimated putative grid orientation and quantified hexadirectional modulation using a cross-validated approach in which orientation was estimated using data from two functional runs and tested on independent data (see Methods). Examining signals in a region of vmPFC previously identified to exhibit six-fold symmetry [3], we identified trials with trajectories aligned to the estimated grid axes and computed multivoxel pattern similarity across trial pairs (Fig. 2b, c). Consistent with grid-like coding, response patterns in portions of vmPFC (Fig. 2a) were more similar for trajectory pairs aligned with the same 60° periodic grid orientation than for those offset by 30° (Δ*r* = 0.0072, 95% CI [0.0034, 0.0112], *d* = 0.6125, *p* = .0001; Fig. 2d, f). This effect was specific to six-fold periodicity characteristic of grid-like coding (six-fold vs. other folds: Δ*r* = 0.0088, 95% CI [0.0046, 0.0133], *d* = 0.9223, *p* = .0005; Fig. 2e). Assessing robustness to coordinate choice, we observed evidence of six-fold modulation at other coordinates in vmPFC reported in past studies [3,5] (S3 Table) and in a searchlight analysis (S6-S7 Fig and S4 Table).

**Figure 2.**
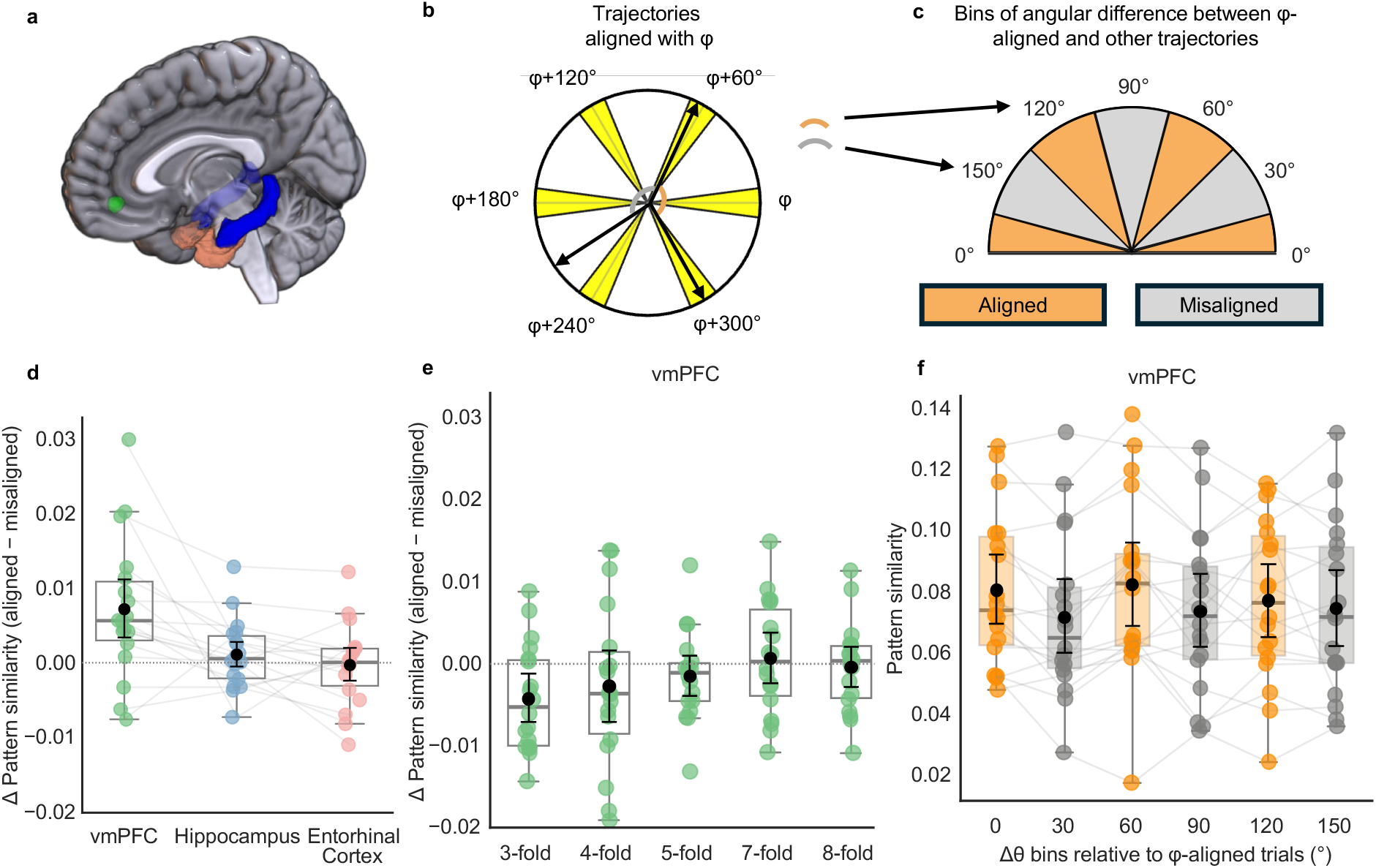
Grid-like coding of affective space in ventromedial prefrontal cortex. **a**, Masks for regions of interest: ventromedial prefrontal cortex (vmPFC; green), hippocampus (blue), and entorhinal cortex (pink). **b**, Trajectories with angles within ± 7.5° to the estimated grid orientation φ (mod 60°) were defined as φ-aligned (yellow). **c**, Trials were classified as aligned (within 0° ± 15°, mod 60°; orange) or misaligned (within 30° ± 15°, mod 60°; gray) to the φ-aligned trials. Two example trial pairs were shown for the 120° and 150° bins (see panel b). **d**, Pattern similarity differences between aligned-aligned and aligned-misaligned trial pairs across regions. **e**, Effects for control periodicities in vmPFC (three-fold: Δ*r* = -0.0043, 95% CI [-0.0068, -0.0017], *d* = -0.5420, *p* = .9989; four-fold: Δ*r* = -0.0028, 95% CI [-0.0064, 0.0010], *d* = -0.2451, *p* = .9244; five-fold: Δ*r* = -0.0015, 95% CI [-0.0040, 0.0011], *d* = -0.1986, *p* = .8802; seven-fold: Δ*r* = 0.0007, 95% CI [-0.0025, 0.0041], *d* = 0.0651, *p* = .3592; eight-fold: Δ*r* = -0.0004, 95% CI [-0.0031, 0.0021], *d* = - 0.0558, *p* = .6333). **f**, Pattern similarity as a function of angular bin (colors as in **c**) in vmPFC. Boxplots show the median and interquartile range; whiskers extend to 1.5× the interquartile range. Black circles and error bars denote the mean and 95% confidence interval. Each point represents one participant (*n* = 18 independent participants) and points from the same participants are connected by gray lines. A dotted line marks zero (no aligned-misaligned difference) in panels d and e. vmPFC = ventromedial prefrontal cortex.

No comparable grid-like code was observed in the hippocampus (Δ*r* = 0.0011, 95% CI [-0.0005, 0.0028], *d* = 0.2220, *p* = .0977; Fig. 2d), and six-fold modulation was weaker than that in vmPFC (Δ*r*_hippocampus –_ Δ*r*_vmPFC_ = -0.0060, 95% CI [-0.0105, -0.0020], *d* = -0.6577, *p* = .0042). We also did not observe grid-like responses in the entorhinal cortex (Δ*r* = -0.0002, 95% CI [-0.0024, 0.0021], *d* = -0.0354, *p* = .5860; Fig. 2d). Temporal signal-to-noise and mean signal in this region were lower (S8 Fig), which may have limited sensitivity to detect subtle effects. To account for potential confounds that could increase pattern similarity between aligned trials, including category, face identity, phrase context, or proximity in valence and arousal (S9 Fig), we conducted a control analysis incorporating these variables as nuisance regressors. The six-fold modulation in vmPFC remained robust in this analysis (Δ*r* = 0.0063, 95% CI [0.0025, 0.0106], *d* = 0.77226, *p* = .0020; six-fold vs. other folds: Δ*r* = 0.0075, 95% CI [0.0035, 0.0118], *d* = 0.8303, *p* = .0012; S10 Fig).

To determine whether the hexadirectional effect in vmPFC was robust across different methods for estimating affective space, we repeated the analysis using trajectory angles derived from multidimensional scaling of emotion category ratings (S5 Fig). The grid-like effect in vmPFC was replicated using this alternative geometry (Δ*r* = 0.0051, 95% CI [0.0009, 0.0097], *d* = 0.3810, *p* = .0155; six-fold vs. other folds: Δ*r* = 0.0049, 95% CI [0.0005, 0.0095], *d* = 0.5075, *p* = .0245). Notably, hexadirectional modulation was present in vmPFC when affective space was defined directly from self-reported valence and arousal and when it was estimated using data-driven multidimensional scaling, indicating that grid-like representation in vmPFC is robust to independent methods of defining affective space.

### Generalization of grid-like codes across facial and linguistic stimuli

Dimensional theories of affect propose that a common cognitive structure underlies judgements about the meaning of words, facial expressions, and subjective experience more generally [9]. If vmPFC implements a grid-like basis for affective space, its orientation should generalize across stimuli that differ in perceptual features but have similar meaning and underlying geometry. Estimating grid orientation from behavioral ratings of face stimuli and testing on phrases revealed robust six-fold modulation in vmPFC (Face → Phrase: Δ*r* = 0.0072, 95% CI [0.0013, 0.0135], *d* = 0.5495, *p* = .0196). The reciprocal analysis produced comparable results (Phrase → Face: Δ*r* = 0.0117, 95% CI [0.0013, 0.0224], *d* = 0.5061, *p* = .0234). These cross-modal results suggest that hexadirectional coding in vmPFC reflects a modality-invariant representation of affective space that instantiates the geometric structure described by dimensional models of emotion.

### Grid-like coding in vmPFC is distinct from decision-related signals

The medial prefrontal cortex is widely implicated in encoding decision-related variables [24], such as subjective value [25–27] and confidence [28,29]. To determine whether the observed grid-like signal in vmPFC reflects dynamic decision processes as opposed to a stable representational basis, we examined how grid-like signals relate to behavioral judgments made during scanning. The degree of hexadirectional modulation did not differ between trials in which judgements were consistent and inconsistent with the group consensus in post-scan ratings (Δ*r*_consistent_ *-* Δ*r*_inconsistent_ = 0.0049, 95% CI [-0.0031, 0.0126], *d* = 0.2880, *p* = .1220; Fig. 3b). This finding suggests that the grid-like code was stable across trials rather than dynamically constructed for individual choices.

**Figure 3.**
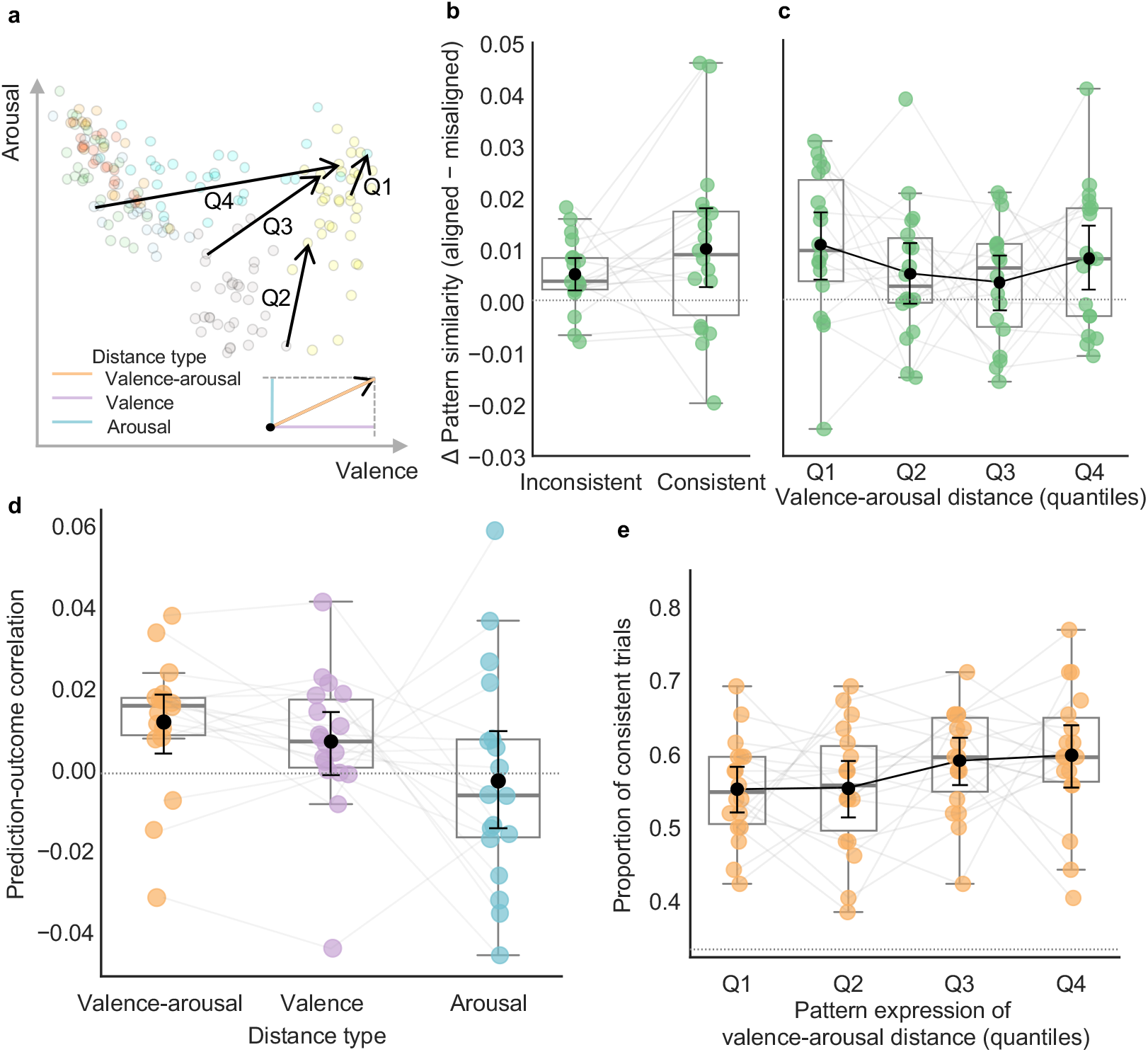
Hexadirectional modulation in vmPFC is distinct from decision-related signals. **a**, Schematic illustration of distance measures in affective space. The background scatter plot shows group-average valence and arousal ratings for all stimuli, as shown in Fig. 1c. Four black vectors illustrate affective trajectories spanning different quantiles of the two-dimensional Euclidean distance. Three types of distance are defined for the transition between a starting stimulus (black dot) and ending stimulus (black arrowhead), including the two-dimensional Euclidean distance (orange), and its one-dimensional projections onto the valence (purple) and arousal (blue) axes. **b**, Pattern similarity differences (aligned-aligned vs. aligned-misaligned) for trial pairs in which both trials had matching in-scanner choices and post-scan ratings (‘consistent’), versus pairs containing at least one mismatch (‘inconsistent’). **c**, Pattern similarity differences in vmPFC across quartiles of the summed Euclidean distances of the two trials in each pair. **d**, Performance of encoding models trained on Euclidean distance and unidimensional valence and arousal distances to predict fMRI signal in vmPFC. The dotted horizontal line indicates chance-level performance. **e**, Proportion of consistent trials across quartiles of pattern expression of Euclidean distance in vmPFC. The dotted horizontal line indicates chance level. Boxplots show the median and interquartile range; whiskers extend to 1.5 × the interquartile range. Black circles and error bars denote the mean and 95% confidence interval. Each point represents one participant (*n* = 18 independent participants) and points from the same participants are connected by gray lines.

Given the strong relationship between trajectory distance in affective space and behavioral consistency (Fig. 1d), we reasoned that vmPFC responses might encode affective distance, potentially derived from readouts of grid-like codes. We therefore fit multivariate encoding models using leave-one-participant-out cross-validation, predicting vmPFC responses from distance in affective space (Fig. 3a). Models based on the Euclidean distance in a two-dimensional valence-arousal space predicted vmPFC activity (*r* = 0.0127, 95% CI [0.0050, 0.0195], *d* = 0.7994, 38.09% of the noise ceiling, *p* = .0021), and outperformed models based on valence or arousal alone (Δ*r* = 0.0097, 95% CI [0.0054, 0.0141], *d* = 1.0296, *p* = .0001; Fig. 3d).

If vmPFC representations of affective distance guide judgements, their expression should predict behavioral agreement across participants. To test this prediction, we computed trial-wise pattern expression scores by projecting the multivoxel activity onto weights from the distance encoding model. Greater expression of affective distance in vmPFC was associated with more consistent behavioral ratings (*β* = 0.0788, SE = 0.0330, *z* = 2.388, 95% CI [0.0156, 0.1430], *p* = .0169, mixed-effect logistic regression; Fig. 3e). This relationship remained after controlling for stimulus modality (*β* = 0.1048, SE = 0.0532, *z* = 1.969, 95% CI [0.0029, 0.2089], *p* = .0489) and did not differ between face and phrase trials (*β* = -0.0523, SE = 0.0644, *z* = -0.812, 95% CI [-0.1786, 0.0754], *p* = .4169), indicating that the association between distance-related activity and behavior did not depend on stimulus type. Moreover, hexadirectional modulation did not vary with trajectory distance (alignment × distance interaction: *β* = -0.0006, 95% CI [-0.0018, 0.0005], *d* = -0.2384, *p* = .8327; Fig. 3c), indicating that the distance-related signal associated with behavior is not a simple consequence of stronger grid-like patterning. Together, these findings indicate vmPFC maintains a stable grid-like representation of affective space and expresses distance-related signals associated with behavioral judgments.

## Discussion

Examining brain activity as participants tracked transitions in facial expressions and language, we observed hexadirectional modulation in vmPFC consistent with grid-like coding of a two-dimensional affective space organized by valence and arousal. This representation generalized across stimulus modalities and was stable across behavioral responses. These findings link the cognitive structure described in dimensional models of emotion to grid-like coding principles previously identified in spatial and conceptual navigation.

Contemporary theories of emotion emphasize coordination among distributed brain systems, but do not specify the format in which affect is represented. Psychological constructionist accounts emphasize how systems involved in visceromotor control, interoception, and conceptualization interact with one another [30–32]. In such accounts, emotional experiences emerge from the integration of bodily states with contextual representations of the current situation to predict the outcome of possible actions, with vmPFC being one site of integration [33]. Neuroimaging and lesion studies [34] have revealed vmPFC’s involvement in affective self-report [35–37], interpretation of social cues [38,39], and prospective valuation [40,41]. The present findings extend this literature by revealing grid-like coding for affective space in vmPFC, providing a format for organizing emotion knowledge. In this framework, principles implicated in spatial and conceptual navigation may also structure how internal affective states are represented and used to guide behavior. Future work using single-unit recordings will be important to determine whether such grid-like population codes reflect periodic tuning at the level of individual neurons, as observed in rodent entorhinal grid cells during spatial navigation [42].

Our findings indicate that vmPFC exhibits grid-like coding of an affective space that is largely shared across individuals. This result is consistent with the view that emotion knowledge reflects learned regularities in affective experiences accumulated over time, rather than being constructed anew on each occasion. Responses in vmPFC were less well captured by individual post-scan ratings (S11 Fig), which may reflect variability in scale use and reduced sensitivity to grid-like structure. Importantly, evidence of a shared structure does not imply that emotion concepts are organized identically across individuals. Prior work demonstrates meaningful variation in affective organization related to development [43] and mental health [44], including differences in emotional granularity, or the degree to which individuals differentiate emotion concepts within affective space [45]. If grid-like codes in vmPFC result from the statistical regularities of emotion transitions, then systematic variation in affect dynamics across experiences may give rise to individual differences in hexagonal modulation. Characterizing how affective space and its grid-like basis are shaped by experience will be critical for understanding the development of the affective circumplex and the mechanisms by which the brain builds structured knowledge of the world.

Differing from past studies involving learning [3,5,46], we found that the magnitude of grid-like modulation did not vary with behavioral choices, which were better predicted by distance-related signals in vmPFC. This dissociation dovetails with electrophysiological recordings in non-human primates showing temporally distinct grid-like and choice-related codes in vmPFC [22]. These findings suggest that affective information may be encoded both as abstract coordinate structure and as context-sensitive evaluative signals related to state value [47,48]. In this view, vmPFC maintains a stable grid-like scaffold for affective space that may support decision-relevant computations to meet the needs of the moment [49]. Although the present findings are based on fMRI measures and do not establish the necessity of vmPFC for organizing affective space, perturbation approaches and longitudinal designs can be used to determine how grid-like coding develops and whether it is required for maintaining affective geometry or decision-making.

In sum, we show that vmPFC BOLD signal exhibits hexadirectional modulation as a function of the direction of transitions between emotion concepts. These findings suggest that the structure evident in self-report may reflect how the brain organizes emotion knowledge using mechanisms similar to those implicated in spatial and conceptual navigation. By identifying a grid-like representational format for affective space, the present work moves the study of emotion beyond the localization of function or predictive modeling toward identifying the processes that may structure affective representations in the human brain.

## Materials and Methods

### Participants

We recruited 20 participants from the Atlanta, GA area. Participants were healthy adults fluent in English, with normal or corrected-to-normal vision, and no contraindications for MRI. The study was approved by the Institutional Research Board at Emory University, and all participants provided informed consent prior to participation. Data from 18 participants (13 females; age: mean = 39.56, SD = 13.19, range = 21-60 years) were analyzed. One participant withdrew due to claustrophobia, and fMRI data from one participant were excluded due to excessive head motion (maximum framewise displacement > 10 mm). Another participant did not perform the post-scan rating task and was therefore excluded from analyses using individual ratings.

### Experimental paradigm

#### Stimuli

Photos of facial expressions were selected from the Karolinska Directed Emotional Faces database [50]. This set of photos depicts facial expressions of human models, and represents seven categories of emotions: happy, surprised, angry, afraid, sad, disgusted, and neutral. A total of 78 photos from 26 identities were used in this experiment. To construct trajectories through affective space, pairs of facial expressions from different emotion categories and the same identity were morphed into dynamic transitions using the Abrosoft FantaMorph 5 software.

Linguistic stimuli (including both multi-word expressions and single words) were selected from eight everyday settings, including ‘shopping’, ‘park’, ‘forest’, ‘school’, ‘food’, ‘animal’, ‘weather’, and ‘social’, chosen to provide distinct semantic contexts in which the same emotion categories can be conveyed. For each setting, a large candidate pool of phrases was generated using ChatGPT-4o with the following prompt ‘create a list of nouns related to [the current setting] and related to the following emotions: surprise, sadness, joy, fear, disgust, anger, neutral’. Valence, arousal, and associated emotion category for all candidate phrases were validated in a pilot study with 10 participants recruited from the Department of Psychology at Emory University. The final set of stimuli was screened such that the highest-rated category for each item matched the category assigned during generation and had a mean rank < 1.5 across pilot participants. A total of 126 phrases were used in this experiment. Pilot phrase ratings were used to guide stimulus selection and were not included in any analyses reported here.

Valence and arousal ratings used to inform the selection of the trajectories come from normative ratings of the face stimuli [51] and the averaged ratings across the pilot participants for phrases. We selected 84 face trajectories and 124 sequences of phrases for use in the scanner, aiming for balanced coverage across emotion categories, settings, and trajectory angles. For descriptive purposes, we visualized how stimulus properties varied across trajectory angles using data from the current study in S9 Fig. To formally test whether any of these factors could account for the observed hexadirectional modulation, category, context (identity or setting), and valence and arousal differences were included as nuisance regressors in control analyses (see Methods, ‘Hexadirectional modulation analysis’).

#### fMRI paradigm

Participants completed an affective judgment task with concurrent fMRI. In this task, participants made judgments about facial expressions and sequences of phrases. The scanning session was divided into four runs, with two runs involving stimuli from either modality. Stimuli were presented using PsychoPy v2022.2.5 [52]. The order of the two modalities (faces and phrases) and the order of the two runs within each modality were counterbalanced across participants. We performed block-wise pseudo-randomization of the order of trials within each run. For facial expressions, trajectories from the same identity were presented in a block structure; for phrases, trajectories from the same setting were grouped, and participants were cued to the current setting at the beginning of each block. Blocks lasted on average 4.73 trials (range = 1-18, SD = 5.48). Block length varied from 2.33 trials (range = 1-9, SD = 1.97) for facial expressions and 15.5 trials (range = 14-18, SD = 1.69) for phrases. This grouping was used to reduce perceptual and semantic variability across consecutive trials and to encourage participants to track affective transitions within a stable context.

In each trial of face runs, participants were presented with a 5-s morph of facial expressions, starting with one of the seven emotion categories and ending with a different category. Similarly, the stimulus in each trial of the phrase runs consisted of a starting phrase (2.5 s) followed by an ending phrase (2.5 s), forming a transition between two different emotion categories. In both the face and phrase runs, each stimulus was followed by a 1-to-5 s jittered fixation cross (mean = 2 s). After fixation, participants had 3 s to indicate whether the valence or arousal of the stimulus increased, decreased, or stayed the same. Participants made responses using their index and middle fingers on a button box in the right hand. Visual feedback was provided with a red dot to indicate ‘no change’, an ‘increase’, or a ‘decrease’. To disentangle the movement of the dot from the movement in the valence-arousal space, both ‘increase’ and ‘decrease’ were designed to be on the right of the ‘no change’. Each trial probed only one dimension (valence or arousal), with trials evenly split across dimensions. Participants were asked to track changes in both valence and arousal throughout the task, as the probed dimension was revealed at the start of the response period.

#### Post-scan self-report ratings

After the scan, participants rated valence, arousal, and seven emotion categories on individual face expressions and phrases. The scales for valence and arousal were presented on one screen, and they ranged from ‘extremely unpleasant/calm’ to ‘extremely pleasant/activated’. The scales for the seven emotion categories (happy, surprised, angry, afraid, sad, disgusted, and neutral) were presented on another screen following the valence-arousal ratings. Category ratings ranged from ‘not at all [category name]’ to ‘extremely [category name]’. All scales were continuous and ticks were provided as anchor points for participants to more easily make the ratings. Participants used a mouse to click on the scale to make the ratings in a self-paced way.

### Behavioral data analysis

#### Low-dimensional organization of emotion concepts

To assess whether emotion concepts are organized in a two-dimensional affective space in the self-report data, we performed multidimensional scaling (MDS) on the group-average ratings of emotion categories. For each stimulus, ratings on the seven emotion categories were treated as a vector, and pairwise cosine distances were computed between all stimulus pairs. The resulting dissimilarity matrix was submitted to metric MDS with dimensionality varied from 1 to 5. Model fit was evaluated using Kruskal stress [53]. The optimal dimensionality was determined based on the elbow in the scree plot of stress values.

We used Procrustes analysis [54] to align the two-dimensional MDS solution to the valence-arousal space derived from direct ratings. To assess the statistical significance of this alignment, we performed a permutation test in which the correspondence between stimuli in the valence–arousal space and the category-derived MDS space was randomly shuffled. The null distribution of Procrustes disparities was generated from 10,000 permutations, and a one-sided *p*-value was computed as the proportion of permuted Procrustes disparities less than or equal to the observed disparity.

#### Cross-modal similarity of affective structure

To test whether the affective structure revealed in self-report is similar across stimulus modalities, we constructed separate dissimilarity matrices for face and phrase stimuli. For valence-arousal ratings, Euclidean distances were computed between stimulus pairs in the two-dimensional valence-arousal space. For emotion category ratings, cosine distances were computed between category-rating vectors. Because face and phrase stimulus sets lacked one-to-one correspondence, we used the Gromov-Wasserstein (GW) distance [55] to quantify the similarity between their structures. The Conditional Gradient algorithm [56] was used (as implemented in the Python Optimal Transport library) to estimate an optimal transport plan that aligns the two structures by minimizing the expected squared difference between their pairwise distances under the alignment. Before entering the GW computation, valence and arousal ratings were z-scored within each modality to make scales of the two spaces comparable. To test statistical significance, we performed permutation tests by disrupting the affective structure obtained from face stimuli (i.e., independently shuffling valence and arousal values or category rating vectors across stimuli), while holding the phrase-stimulus structure fixed. The null distributions of GW distances were generated from 10,000 permutations, and one-sided *p*-values were computed as the proportion of permuted GW distances that were less than or equal to the observed distance.

#### Behavioral sensitivity to affective distance during the fMRI task

To validate that participants’ in-scanner judgments reflected the same evaluation of the stimuli captured by their post-scan ratings, we tested whether the two measures were consistent across trials. Trials were labeled as being consistent when participants’ in-scanner choice matched the difference in post-scan ratings of the judged dimension between the ending and starting stimuli. Specifically, an absolute rating difference smaller than 1 (i.e., the distance between two adjacent anchor points on the rating scale) was considered consistent with a ‘no change’ choice. If the difference was higher than 1, a choice of ‘increase’ in the scanner was considered to be consistent and vice versa if the difference was lower than -1. We then used a mixed-effects logistic regression to model the probability of a consistent choice (coded as 1) as a function of the post-scan rating change, with random intercepts and slopes for each participant.

### MRI data acquisition

Gradient-echo echo-planar imaging (EPI) data were acquired on a 3T Siemens MAGNETOM Prisma scanner with 32-channel parallel imaging located at the Facility for Education and Research in Neuroscience at Emory University. T2*-weighted functional data were acquired using the following parameters: repetition time (TR) = 1250 ms; echo times (TEs) = 15 ms / 37.16 ms / 59.3 ms; flip angle = 55°; number of slices = 56 interleaved; voxel size = 2.5 mm isotropic; field of view = 210 mm; multiband acceleration factor = 4; bandwidth = 2381 Hz/pixel. The slice angle was set to 10° relative to the anterior-posterior commissure line to reduce signal loss in basal frontal regions [57,58]. A pair of fieldmap scans with reversed phase-encoding directions (posterior-anterior and anterior-posterior) was acquired to enable correction of susceptibility-induced distortions in the functional images. Each fieldmap acquisition used the same voxel size, slice coverage and field of view as the functional scans, with TR = 7220 ms, TE = 73 ms, and flip angle = 90°.

To facilitate normalization of EPIs to standard space, structural images were acquired using T1 MPRAGE: TR = 1900 ms; TE = 2.26 ms; flip angle = 9°; number of slices = 192; voxel size = 1 mm isotropic; field of view = 256 mm × 256 mm; GRAPPA acceleration factor = 2; bandwidth = 199 Hz/pixel.

### MRI preprocessing

Results included in this manuscript come from preprocessing performed using *fMRIPrep* 25.0.0 [59,60] (RRID:SCR_016216), which is based on *Nipype* 1.9.2 [61,62] (RRID:SCR_002502).

#### Preprocessing of B0 inhomogeneity mappings

One set of fieldmaps was found available within the input BIDS structure for each subject. A *B0*-nonuniformity map (or *fieldmap*) was estimated based on two (or more) echo-planar imaging (EPI) references with FSL’s topup [63].

#### Anatomical data preprocessing

A total of 1 T1-weighted (T1w) images were found within the input BIDS dataset. The T1w image was corrected for intensity non-uniformity (INU) with N4BiasFieldCorrection [64], distributed with ANTs 2.5.4 [65] (RRID:SCR_004757), and used as T1w-reference throughout the workflow. The T1w-reference was then skull-stripped with a *Nipype* implementation of the antsBrainExtraction.sh workflow (from ANTs), using OASIS30ANTs as target template. Brain tissue segmentation of cerebrospinal fluid (CSF), white-matter (WM) and gray-matter (GM) was performed on the brain-extracted T1w using FSL’s fast [66] (RRID:SCR_002823). Volume-based spatial normalization to one standard space (MNI152NLin2009cAsym) was performed through nonlinear registration with antsRegistration (ANTs 2.5.4), using brain-extracted versions of both T1w reference and the T1w template. The following template was selected for spatial normalization and accessed with *TemplateFlow* [67] (24.2.2): *ICBM 152 Nonlinear Asymmetrical template version 2009c* [68] (RRID:SCR_008796; TemplateFlow ID: MNI152NLin2009cAsym).

#### Functional data preprocessing

For each of the 4 BOLD runs found per subject (across all tasks and sessions), the following preprocessing was performed. First, a reference volume was generated from the shortest echo of the BOLD run, using a custom methodology of *fMRIPrep*, for use in head motion correction. Head-motion parameters with respect to the BOLD reference (transformation matrices, and six corresponding rotation and translation parameters) are estimated before any spatiotemporal filtering using FSL’s mcflirt [69]. The estimated *fieldmap* was then aligned with rigid-registration to the target EPI (echo-planar imaging) reference run. The field coefficients were mapped on to the reference EPI using the transform. The BOLD reference was then co-registered to the T1w reference using mri_coreg (FreeSurfer) followed by FSL’s flirt [70] with the boundary-based registration [71] cost-function. Co-registration was configured with six degrees of freedom.

Several confounding time-series were calculated based on the *preprocessed BOLD*: framewise displacement (FD), DVARS and three region-wise global signals. FD was computed using two formulations following Power [72] (absolute sum of relative motions) and Jenkinson [69] (relative root mean square displacement between affines). FD and DVARS are calculated for each functional run, both using their implementations in *Nipype* (following the definitions by Power et al. 2014 [72]). The three global signals are extracted within the CSF, the WM, and the whole-brain masks. Additionally, a set of physiological regressors were extracted to allow for component-based noise correction [73] (*CompCor*). Principal components are estimated after high-pass filtering the *preprocessed BOLD* time-series (using a discrete cosine filter with 128s cut-off) for the two *CompCor* variants: temporal (tCompCor) and anatomical (aCompCor). tCompCor components are then calculated from the top 2% variable voxels within the brain mask. For aCompCor, three probabilistic masks (CSF, WM and combined CSF+WM) are generated in anatomical space. The implementation differs from that of Behzadi et al. [73] in that instead of eroding the masks by 2 pixels on BOLD space, a mask of pixels that likely contain a volume fraction of GM is subtracted from the aCompCor masks. This mask is obtained by thresholding the corresponding partial volume map at 0.05, and it ensures components are not extracted from voxels containing a minimal fraction of GM. Finally, these masks are resampled into BOLD space and binarized by thresholding at 0.99 (as in the original implementation). Components are also calculated separately within the WM and CSF masks. For each CompCor decomposition, the *k* components with the largest singular values are retained, such that the retained components’ time series are sufficient to explain 50 percent of variance across the nuisance mask (CSF, WM, combined, or temporal). The remaining components are dropped from consideration. The head-motion estimates calculated in the correction step were also placed within the corresponding confounds file. The confound time series derived from head motion estimates and global signals were expanded with the inclusion of temporal derivatives and quadratic terms for each [74]. Frames that exceeded a threshold of 0.5 mm FD or 1.5 standardized DVARS were annotated as motion outliers. Additional nuisance timeseries are calculated by means of principal components analysis of the signal found within a thin band (*crown*) of voxels around the edge of the brain, as proposed by Patriat, Reynolds, and Birn (2017) [75]. All resamplings can be performed with *a single interpolation step* by composing all the pertinent transformations (i.e. head-motion transform matrices, susceptibility distortion correction when available, and co-registrations to anatomical and output spaces). Gridded (volumetric) resamplings were performed using nitransforms, configured with cubic B-spline interpolation.

Many internal operations of *fMRIPrep* use *Nilearn* 0.11.1 [76] (RRID:SCR_001362), mostly within the functional processing workflow. For more details of the pipeline, see the section corresponding to workflows in *fMRIPrep*’s documentation (https://fmriprep.readthedocs.io/en/latest/workflows.html).

### Definition of regions of interest

Masks of the hippocampus and entorhinal cortex were obtained from the Julich-Brain Cytoarchitectonic Atlas [77]. The vmPFC mask was obtained by drawing a sphere with radius of 5 mm around the peak voxel of the hexadirectional effect reported by Constantinescu et al. [3] (MNI peak voxel coordinates: [6, 44, -10]; 69 voxels). Additional vmPFC peak coordinates reported in prior studies ([6, 46, -10] [5], [2, 28, -20] [3], and [-8, 42, 0] [5]) were also examined using identical 5-mm-radius spherical regions of interest. For searchlight visualization within vmPFC, a larger anatomical mask including Brodmann areas 10, 11, 14, 24, 25, and 32 was defined based on a multimodal cortical parcellation of Glasser et al. [78].

### General linear models

First-level general linear models were applied to preprocessed fMRI data using SPM 12 software (Wellcome Trust Centre for Neuroimaging, UK). For each participant, a separate model was estimated that included: 1) one regressor per trajectory of facial expressions or phrases, modeled as a 5 s boxcar function, 2) a regressor for button presses, modeled using a stick function, and 3) a 3-s boxcar regressor for the response period. All regressors were convolved with the canonical hemodynamic response function provided in SPM. To account for head motion, 24 nuisance regressors derived from spatial realignment (i.e., three translations, three rotations, and their first- and second-order temporal derivatives) were included, along with constant terms modeling the mean signal for each imaging session.

### Hexadirectional modulation analysis

To test whether voxel-wise activity patterns reflected six-fold symmetry within each region of interest or searchlight, we performed a multivariate hexadirectional modulation analysis using a cross-validated procedure following Bao et al. [5]. For each participant, the data were split into two halves of runs, with one for estimating grid orientation (training set) and the other for testing modulation effects (test set). In the training set, we regressed cos(6θ) and sin(6θ) against single-trial βs estimated for the trajectories. The resulting regression coefficients (β_cos_ and β_sin_) were averaged across voxels within each region. The participant-specific grid orientation φ was then computed as

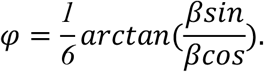

In the test half, trials were classified as *aligned* if their trajectory angle θ was within ± 7.5° of φ (mod 60°). We computed the Pearson correlation between all pairs of response patterns for each region. Correlations were averaged across pairs of trials that were both aligned (aligned-aligned pairs, angular difference within 0° ± 15° mod 60°) and mispatched pairs (aligned-misaligned pairs with an angular difference within 30° ± 15° mod 60°). Comparisons where both trials were misaligned were excluded to avoid low-signal, high-noise pairs [5].

Trajectory angles (θs) used in the analysis were obtained either from ratings of valence and arousal or coordinates in a two-dimensional MDS solution derived from pairwise cosine distances of emotion category ratings. To obtain stable MDS-derived angles, we repeated the MDS procedure with ten random initialization seeds. One solution was selected as the reference, and all remaining solutions were aligned to it using Procrustes analysis. Trajectory angles were then computed separately for the reference and the aligned solutions, and the resulting angles were averaged across the ten MDS solutions to yield estimates of θ.

To quantify the alignment effect while controlling for potential confounds, the following linear model was estimated,

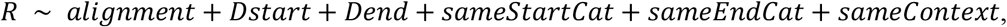

where *R* denotes the pairwise correlation, *alignment* (1 = aligned-aligned, 0 = aligned-misaligned) is the predictor of interest, *Dstart* and *Dend* are Euclidean distances for each trajectory in the affective space, and *sameStartCat, sameEndCat*, and *sameContext* are binary regressors (1 = same, 0 = different) accounting for emotion category and stimulus context (face identity or phrase setting). Because different trajectories were sampled randomly for different levels of context and emotion categories, these additional variables control for effects unrelated to hexagonal modulation.

To assess the specificity of the sixfold modulation, we repeated the same procedures for alternative periodicities of three-, four-, five-, seven-, and eight-fold symmetry, corresponding to angular symmetries spaced every 120° (3-fold), 90° (4-fold), 72° (5-fold), 51.4° (7-fold), and 45° (8-fold). For each periodicity k, φ was estimated using the same procedure as for the sixfold model but applying the corresponding periodic factor. Trials were first classified as aligned if their trajectory angle θ was within ± (360° / (8k)) of φ (mod 360° / k). For each aligned trial, we computed correlations with all other trials and labeled each pair as aligned-aligned or aligned-misaligned depending on whether the angular difference between trajectories was near 0° or half of the symmetry spacing (360° / (2k)).

Statistical significance of group-level effects (both against 0 and for differences between regions or folds) was assessed using sign-flipping with 10,000 iterations. For each iteration, participant-specific aligned-misaligned contrast values or regression coefficients for alignment were randomly sign-inverted to generate a null distribution of group means. One-tailed *p*-values were then computed as the proportion of permuted group means that were greater than or equal to the observed group mean. To estimate uncertainty in group-level effects, we performed nonparametric bootstrap resampling (10,000 iterations) across participants, producing a distribution of group means to derive 95% confidence intervals using the percentile method [79]. Effect size was then calculated by dividing the observed group mean by the standard deviation of the bootstrap distribution, multiplied by the square root of the sample size.

### Distance encoding

We built multivariate encoding models to test whether different distance measures capture variation in BOLD signal in each region of interest. For each region, we used partial least squares regression (SIMPLS algorithm [80]) to predict multivoxel patterns of single-trial β estimates from distances in a cross-validated way. Models were trained using data from all participants except one and tested on held-out participants (leave-one-participant-out cross-validation). Encoding performance was quantified as the Pearson correlation between the predicted and observed multivoxel time series, computed across trials and then averaged across voxels within each region of interest to yield a single performance value per region and participant. To estimate an upper bound on model performance, the noise ceiling was calculated by correlating each held-out participant’s observed time series of brain activity with the mean time series of all other participants. One noise ceiling value was obtained for each region by averaging across held-out participants and voxels. Group-level inference for encoding model performance was conducted using the same sign-flipping and bootstrap resampling procedures as above (see ‘Hexadirectional modulation analysis’).

### Relationship between distance-related brain activity and choice behavior

To test whether the distance-related brain pattern predicts choice behavior in the scanner, we computed a pattern expression score for each trial of the held-out participant. Each participant’s observed multivoxel activity pattern (Y_test_) was projected onto the distance-encoding weights (β_d_) obtained from other participants (Y_test_ ⋅ β_d_), producing one value reflecting how strongly the response pattern on each trial expressed the distance-coding axis. These trial wise pattern expression scores were then entered into a mixed-effects logistic regression model to predict agreement between the judgements made in the scanner and the post-scan ratings (see ‘Behavioral data analysis’ for how consistency measures were obtained). The model included random intercepts and slopes for each participant to account for individual variability. In the control analysis, we tested whether this relationship was driven by lower-level stimulus properties by including stimulus modality (face vs. phrase) and its interaction with the pattern expression score as fixed effects in the model.

All analyses were performed using CanlabCore Tools

(https://github.com/canlab/CanlabCore) and SPM 12

(https://www.fil.ion.ucl.ac.uk/spm/software/spm12/), as well as custom MATLAB (R2024a), python (3.9.18) and R (4.4.1) code.

## Supporting information

SupportingInfo

## >Acknowledgements

We would like to thank Grace Tao for her help with data collection.

