## SupportingInfo for "A grid-like basis for affective space in human ventromedial prefrontal cortex"

### Supporting Figures

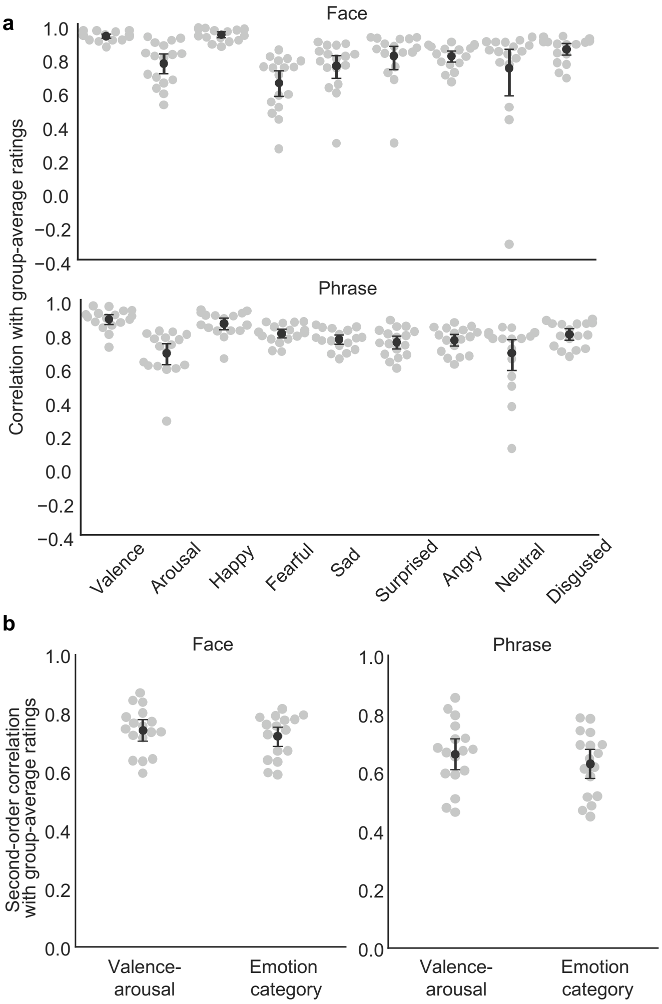


**S1 Fig**. **Agreement between individual and group-average post-scan ratings. a,** First-order agreement. Pearson correlations between individual and group-average ratings for face and phrase stimuli. **b,** Second-order agreement. Spearman correlations between individual and group-average dissimilarity matrices derived from valence-arousal ratings (Euclidean distance) and emotion category ratings (cosine distance) for face and phrase stimuli. Black circles and error bars denote the mean and 95% confidence interval. Each point represents one subject (*n* = 18 independent subjects).

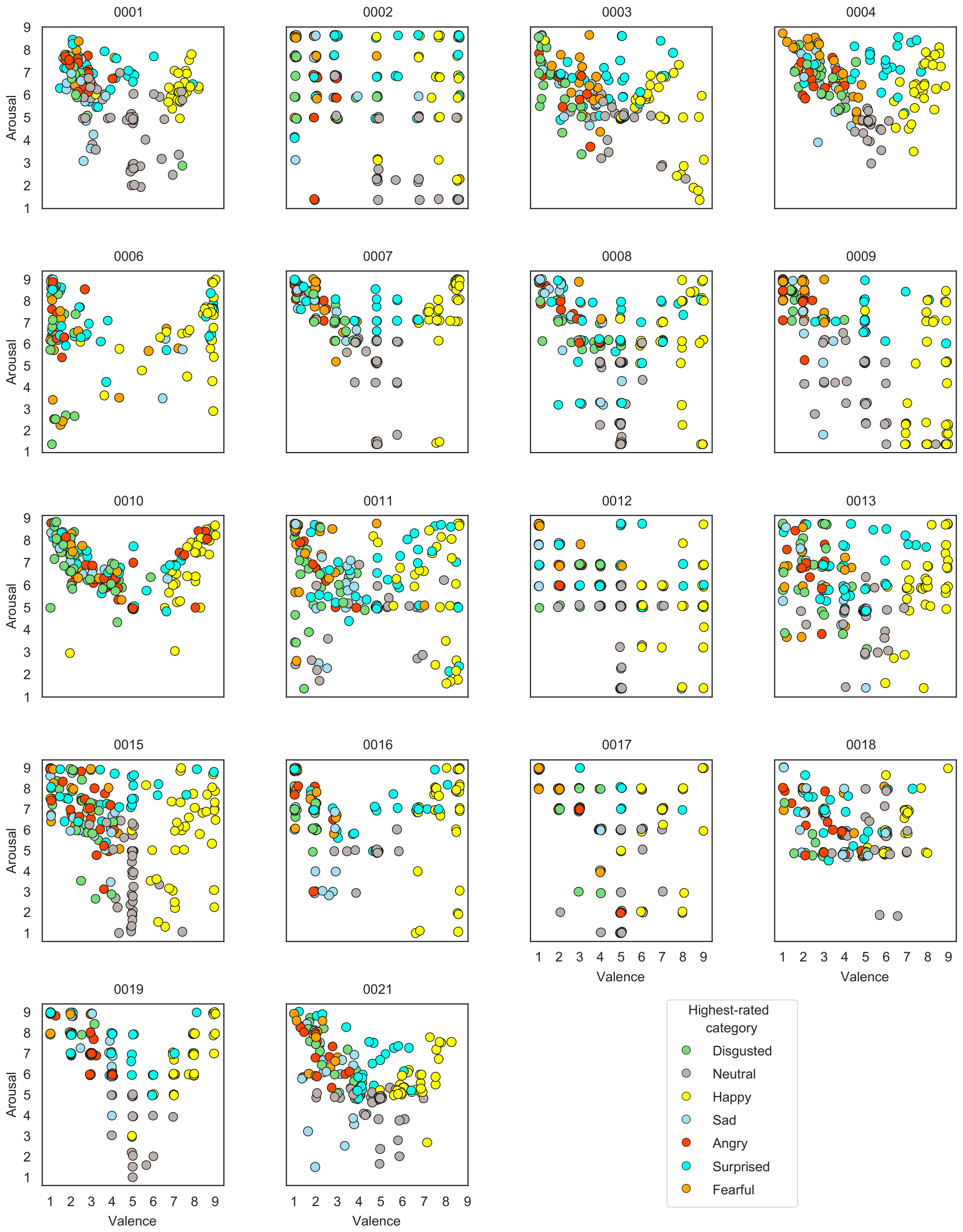


**S2 Fig. Valence and arousal ratings for each participant**. Each subplot shows data from one participant. Each point represents one stimulus (including both faces and phrases) and is colored according to the emotion category with the highest rating.

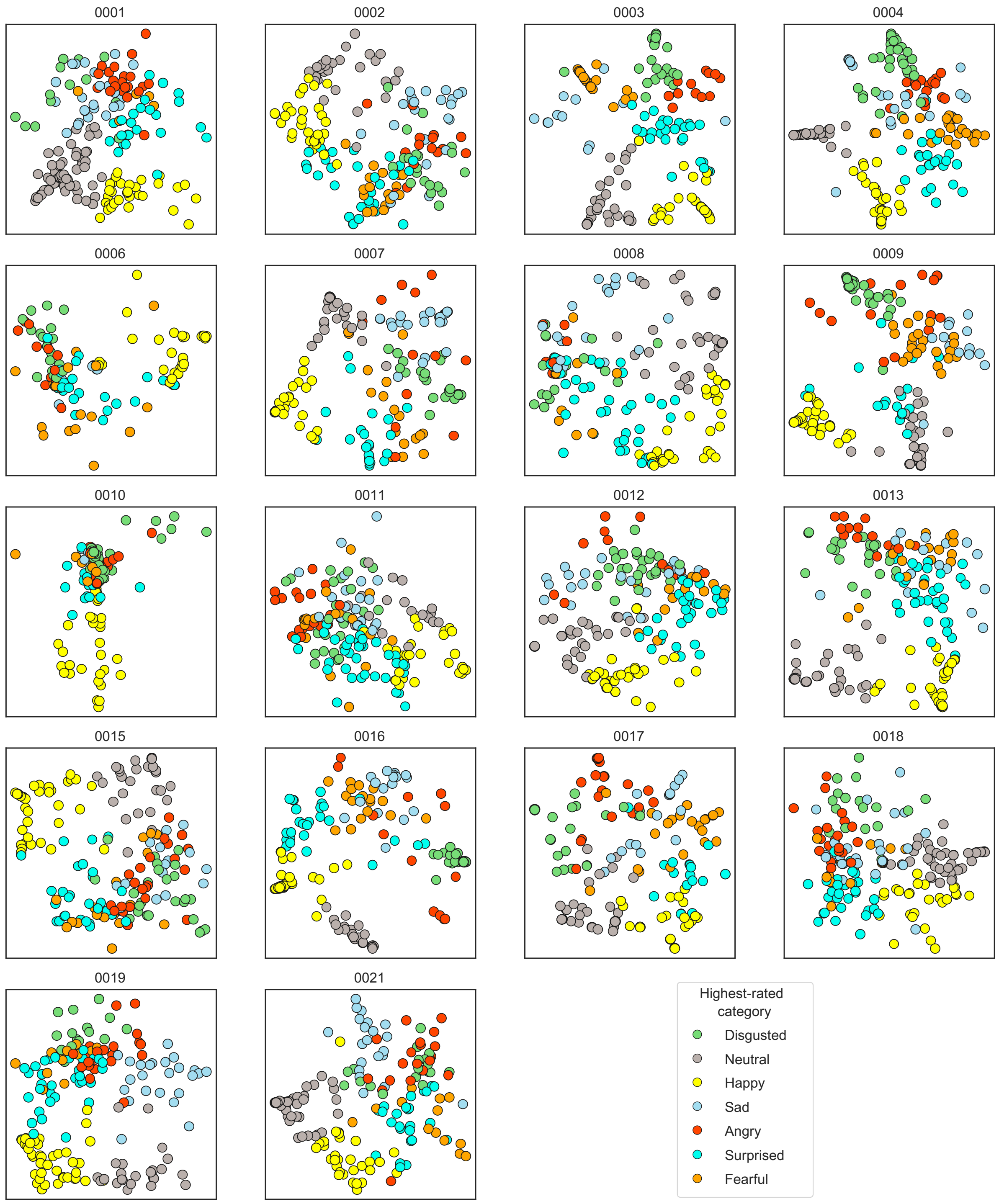


**S3 Fig. Multidimensional scaling (MDS) of emotion category ratings for each participant.** Each subplot shows the two-dimensional MDS solution for data from one participant. Each point represents one stimulus (including both faces and phrases) and is colored according to the emotion category with the highest rating.

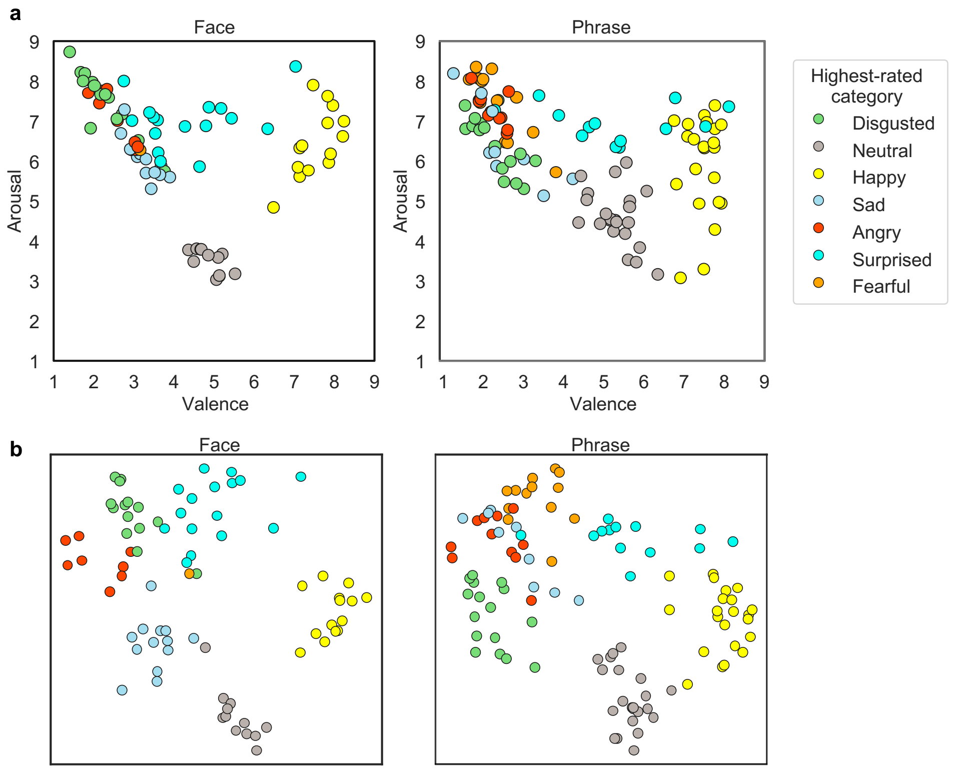


**S4 Fig. Low-dimensional embeddings of emotions conveyed by face and phrase stimuli.** **a,** Group-average valence-arousal ratings of face and phrase stimuli. Faces and phrases exhibited highly similar structures (Gromov-Wasserstein distance = 0.19, permutation $p$ < .0001). **b,** Multidimensional scaling solutions derived from group-average category ratings for face and phrase stimuli, aligned to the valence-arousal space in **a** using Procrustes analysis. Faces and phrases exhibited highly similar structures in category ratings (Gromov-Wasserstein distance = 0.01, permutation $p$ < .0001). Each point represents one stimulus and is colored according to the emotion category with the highest rating.

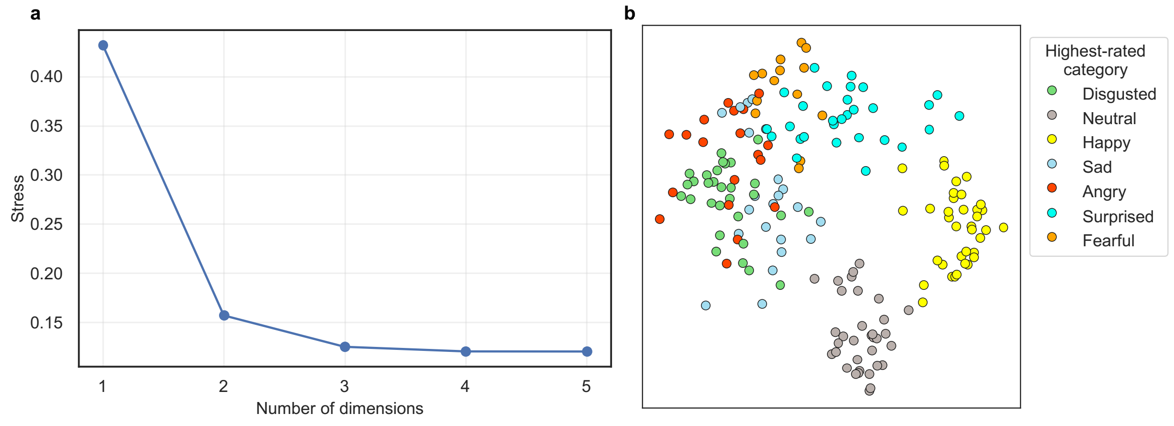


**S5 Fig**. **Multidimensional scaling (MDS) of emotion category ratings. a,** Scree plot of Kruskal stress for MDS solutions on cosine distances between group-average ratings of emotion categories. Note the elbow at two dimensions (Kruskal stress = 0.16). **b,** Two-dimensional MDS solution aligned to the valence-arousal space (see Fig. 1c in the main text) using Procrustes analysis (disparity = 0.20, permutation $p$ < .0001). Each point represents one stimulus and is colored according to the emotion category with the highest rating.

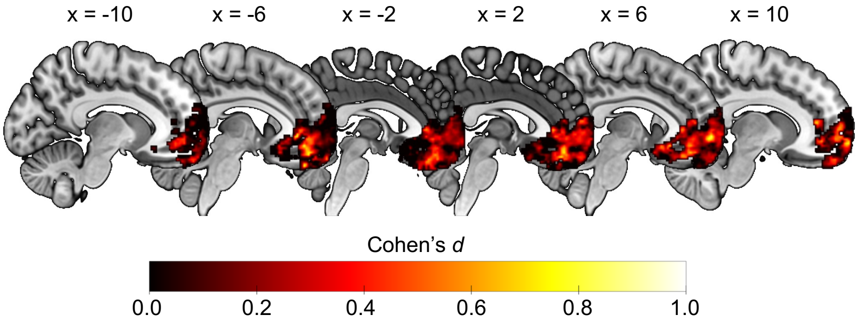


**S6 Fig. Voxelwise effect sizes for searchlight hexadirectional modulation analysis within ventromedial prefrontal cortex.** Searchlight maps (radius = 5 mm) reflect Cohen's *d* values of pattern similarity differences between aligned and misaligned trial pairs across participants ($n$ = 18 independent participants). Warmer colors indicate greater similarity for aligned relative to misaligned trial pairs. The maps are shown within a mask of prefrontal cortex comprising Brodmann areas 10, 11, 14, 24, 25, and 32. Only voxels with positive effect sizes are displayed.

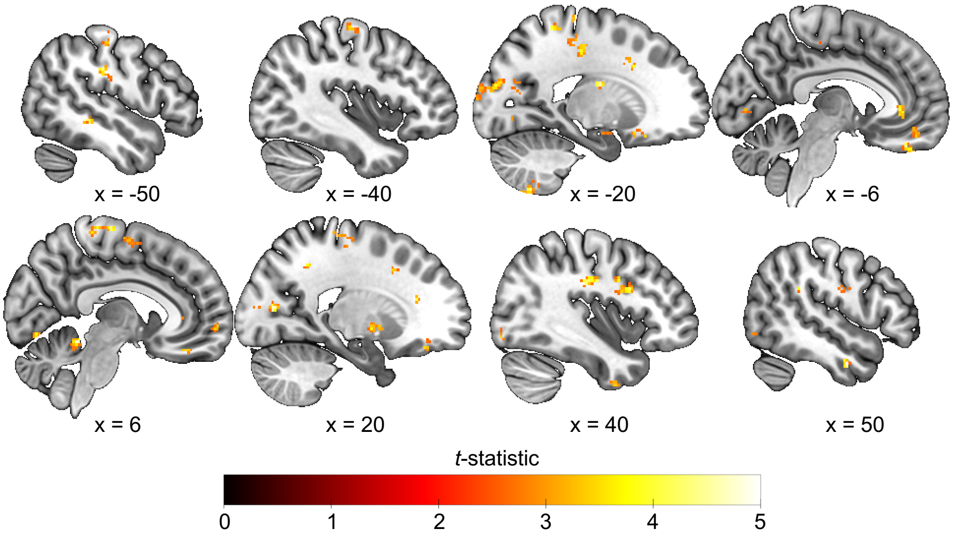


**S7 Fig. Whole-brain searchlight map of hexadirectional modulation.** Parametric $t$-statistic maps show effects of hexadirectional modulation derived from an exploratory whole-brain searchlight analysis (radius = 5 mm). Voxels were thresholded at $p < .01$ (one-tailed; uncorrected), with a minimum cluster extent of $k \geq 20$ contiguous voxels for visualization. Warm colors reflect greater similarity for aligned relative to misaligned trial pairs. No voxels survived false discovery rate correction at a threshold of $q < .05$.

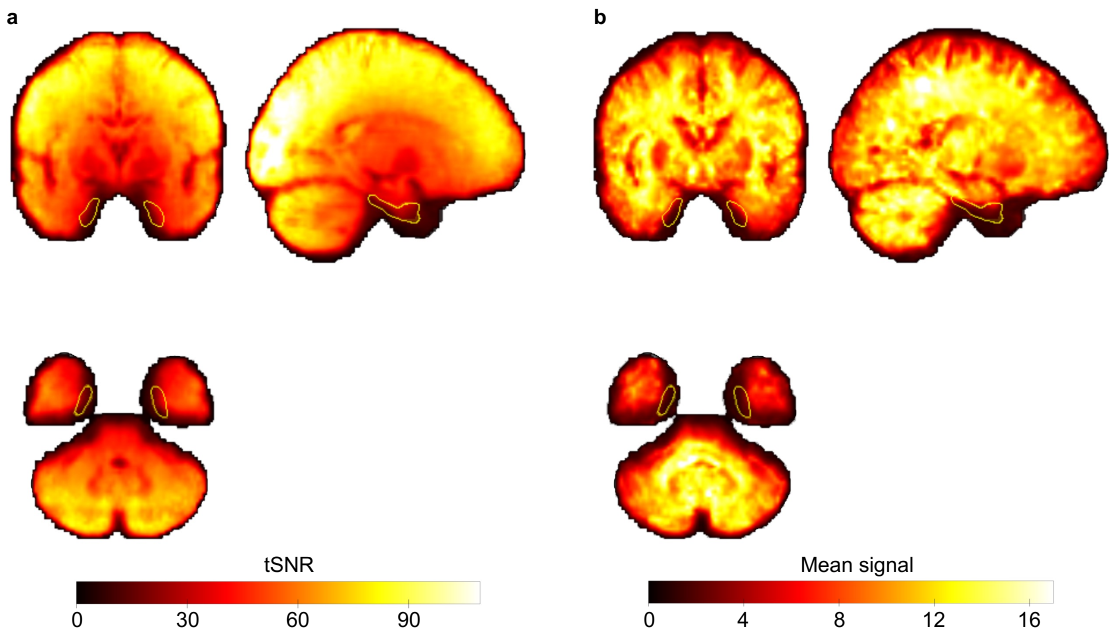


**S8 Fig. Whole-brain maps of signal quality. a,** Group temporal signal-to-noise ratio (tSNR) map. For each participant and run, voxelwise tSNR maps were created as the mean preprocessed BOLD signal divided by its standard deviation over time. The parametric map shows tSNR values averaged across runs and participants ($n$ = 18 independent participants). **b,** Mean BOLD signal across participants. For each participant, voxelwise signal intensity maps were created by averaging preprocessed BOLD timeseries across all runs. The parametric map shows the group mean of signal intensity divided by the standard deviation across participants ($n$ = 18 independent participants). Note that the ventral surface of the brain (including entorhinal cortex outlined in yellow) exhibits lower tSNR and reduced mean signal intensity.

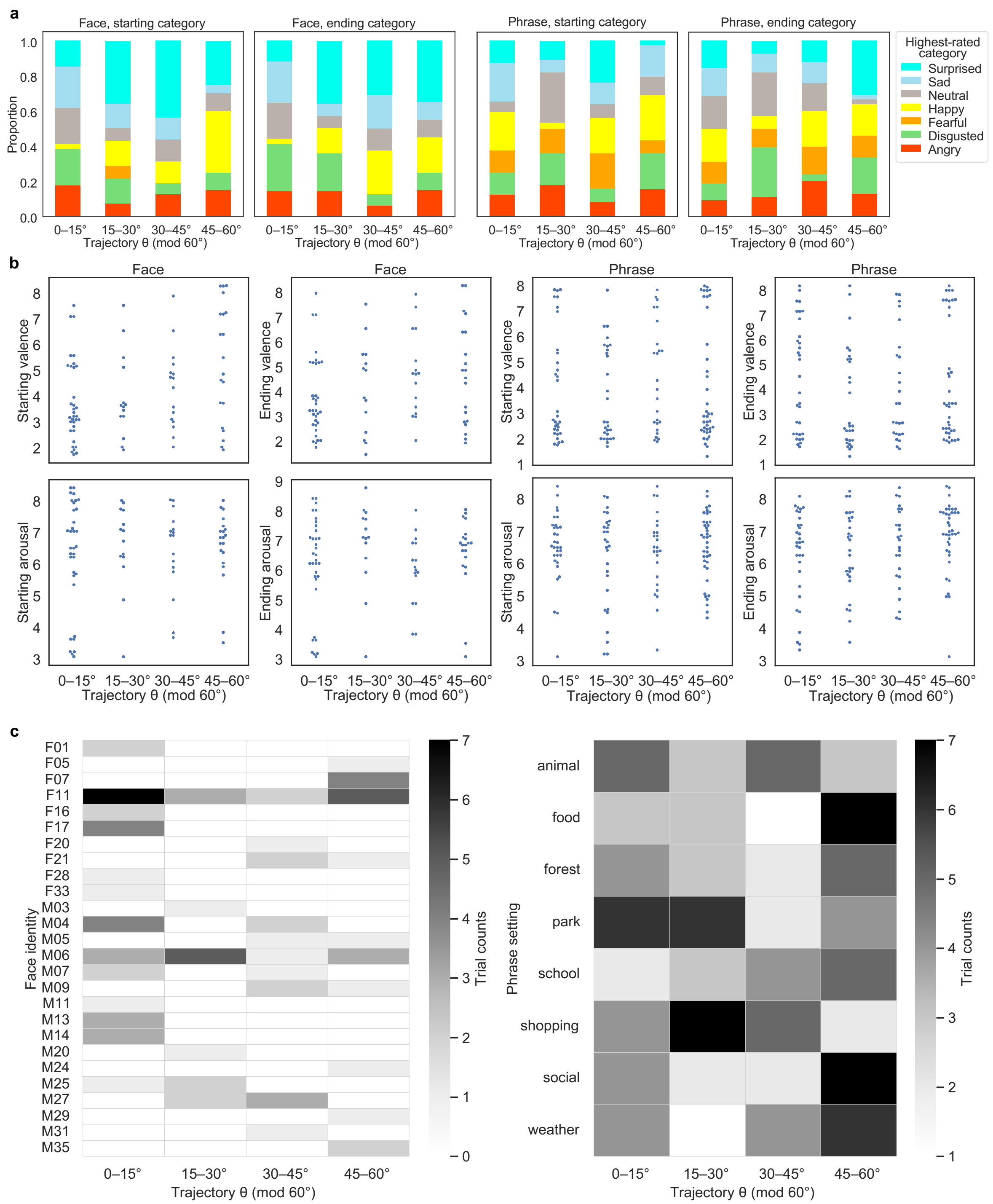


**S9 Fig. Distribution of stimulus properties across trajectory angles. a,** Proportion of highest-rated emotion categories across the four 15° bins of the trajectory angle (θ mod 60°) for starting and ending face and phrase stimuli. Bars show the relative frequency of categories appearing within each bin. **b,** Distributions of valence and arousal ratings across the same four angle bins for starting and ending face and phrase stimuli. Each point represents one stimulus. **c,** Heatmaps showing the number of trials associated with each face identity or phrase setting across the four angle bins.

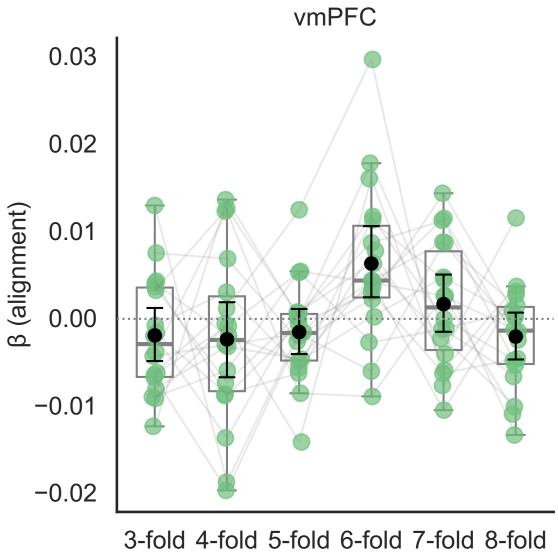


**S10 Fig. Alignment effect across periodicities after controlling for stimulus properties.** Regression coefficients for the alignment effect (aligned-aligned vs. aligned-misaligned trial pairs) are shown across periodicities, estimated from a linear model that controls for distance in valence-arousal ratings, shared emotion categories, and shared face identity or phrase setting. Boxplots show the median and interquartile range; whiskers extend to 1.5× the interquartile range. Black circles and error bars denote the mean and 95% confidence interval. Each point represents one subject ($n$ = 18 independent subjects) and points from the same subjects are connected by gray lines. vmPFC = ventromedial prefrontal cortex.

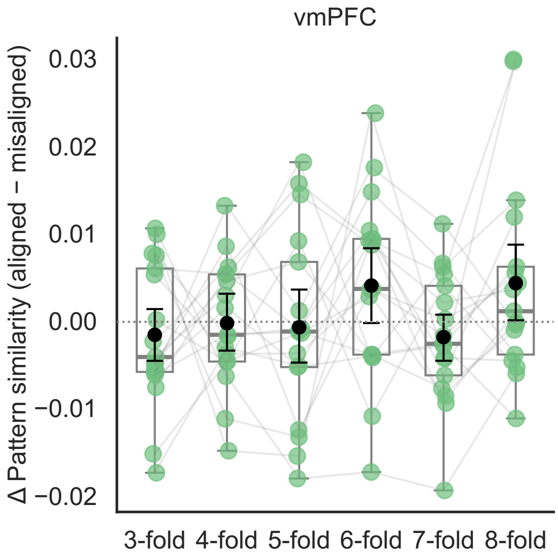


**S11 Fig. Grid-like coding effect in vmPFC using trajectory angles derived from individual ratings.** Each point represents the pattern similarity difference between aligned-aligned and aligned-misaligned trial pairs for one subject ($n$ = 18 independent subjects) and points from the same subjects are connected by gray lines. Boxplots show the median and interquartile range; whiskers extend to 1.5× the interquartile range. Black circles and error bars denote the mean and 95% confidence interval. We did not observe a reliable six-fold effect in vmPFC using trajectory angles obtained from participants' own ratings ($\Delta r$ = 0.0041, 95% CI [-0.0001, 0.0084], $d$ = 0.3294, $p$ = .0617; six-fold vs. other folds: $\Delta r$ = 0.0041, 95% CI [-0.0005, 0.0086], $d$ = 0.4292, $p$ = .0527). vmPFC = ventromedial prefrontal cortex.

**Supporting Tables**

**S1 Table**. **First-order agreement between individual and group-average ratings, and group-level reliability.**

| Item | Modality | Mean *r* | SD of *r* | ICC(2,k) | ICC 95% CI |
|---|---|---|---|---|---|
| Valence | Combined | 0.89 | 0.05 | 0.98 | 0.97, 0.98 |
| Arousal | Combined | 0.7 | 0.14 | 0.92 | 0.90, 0.94 |
| Happy | Combined | 0.88 | 0.05 | 0.97 | 0.96, 0.98 |
| Fearful | Combined | 0.75 | 0.06 | 0.95 | 0.93, 0.96 |
| Sad | Combined | 0.75 | 0.07 | 0.94 | 0.92, 0.95 |
| Surprised | Combined | 0.76 | 0.09 | 0.92 | 0.88, 0.94 |
| Angry | Combined | 0.77 | 0.06 | 0.93 | 0.90, 0.95 |
| Neutral | Combined | 0.7 | 0.21 | 0.92 | 0.90, 0.94 |
| Disgusted | Combined | 0.81 | 0.07 | 0.94 | 0.92, 0.96 |
| Valence | Face | 0.93 | 0.03 | 0.99 | 0.98, 0.99 |
| Arousal | Face | 0.76 | 0.13 | 0.95 | 0.91, 0.98 |
| Happy | Face | 0.93 | 0.04 | 0.99 | 0.98, 0.99 |
| Fearful | Face | 0.65 | 0.16 | 0.89 | 0.80, 0.95 |
| Sad | Face | 0.75 | 0.15 | 0.95 | 0.91, 0.98 |
| Surprised | Face | 0.81 | 0.15 | 0.93 | 0.87, 0.97 |
| Angry | Face | 0.81 | 0.07 | 0.91 | 0.83, 0.96 |
| Neutral | Face | 0.74 | 0.3 | 0.93 | 0.87, 0.97 |
| Disgusted | Face | 0.85 | 0.08 | 0.95 | 0.90, 0.98 |
| Valence | Phrase | 0.88 | 0.06 | 0.98 | 0.97, 0.98 |
| Arousal | Phrase | 0.68 | 0.13 | 0.92 | 0.89, 0.94 |
| Happy | Phrase | 0.86 | 0.07 | 0.97 | 0.96, 0.98 |
| Fearful | Phrase | 0.8 | 0.06 | 0.95 | 0.93, 0.96 |
| Sad | Phrase | 0.76 | 0.06 | 0.93 | 0.91, 0.95 |
| Surprised | Phrase | 0.75 | 0.08 | 0.91 | 0.87, 0.94 |
| Angry | Phrase | 0.76 | 0.08 | 0.93 | 0.90, 0.95 |
| Neutral | Phrase | 0.68 | 0.2 | 0.92 | 0.89, 0.94 |
| Disgusted | Phrase | 0.79 | 0.07 | 0.94 | 0.92, 0.96 |

**S2 Table. Second-order agreement between individual and group-average relational structures.**

| Rating type | Modality | Mean *r* | SD of *r* |
|---|---|---|---|
| Emotion category | Combined | 0.65 | 0.11 |
| Valence-arousal | Combined | 0.67 | 0.1 |
| Emotion category | Face | 0.72 | 0.07 |
| Valence-arousal | Face | 0.74 | 0.08 |
| Emotion category | Phrase | 0.63 | 0.11 |
| Valence-arousal | Phrase | 0.66 | 0.11 |

**S3 Table. Effects of hexadirectional modulation at peak coordinates in ventromedial prefrontal cortex reported in prior conceptual and olfactory navigation studies.**

| Fold | MNI [6, 44, -10] (Constantinescu et al., 2016) | MNI [6, 46, -10] (Bao et al., 2019) | MNI [2, 28, -20] (Constantinescu et al., 2016) | MNI [-8, 42, 0] (Bao et al., 2019) |
|---|---|---|---|---|
| 6 | $\Delta r = 0.0072$, 95% CI [0.0034, 0.0112], $d = 0.6125$, $p = .0001$ | $\Delta r = 0.0055$, 95% CI [0.0014, 0.0096], $d = 0.6207$, $p = .0108$ | $\Delta r = 0.0027$, 95% CI [-0.0001, 0.0056], $d = 0.4339$, $p = 0.0442$ | $\Delta r = 0.0028$, 95% CI [-0.0006, 0.0063], $d = 0.3673$, $p = 0.0734$ |
| 6 vs. other | $\Delta r = 0.0088$, 95% CI [0.0046, 0.0133], $d = 0.9223$, $p = .0005$ | $\Delta r = 0.0064$, 95% CI [0.0016, 0.0113], $d = 0.6025$, $p = .0119$ | $\Delta r = 0.0036$, 95% CI [0.0005, 0.0068], $d = 0.5344$, $p = 0.0191$ | $\Delta r = 0.0045$, 95% CI [0.0014, 0.0078], $d = 0.6539$, $p = 0.0053$ |
| 3 | $\Delta r = -0.0043$, 95% CI [-0.0068, -0.0017], $d = -0.5420$, $p = .9989$ | $\Delta r = -0.0039$, 95% CI [-0.0070, -0.0007], $d = -0.5734$, $p = .9819$ | $\Delta r = -0.0048$, 95% CI [-0.0074, -0.0019], $d = -0.7906$, $p = 0.9967$ | $\Delta r = -0.0019$, 95% CI [-0.0051, 0.0013], $d = -0.2662$, $p = 0.8557$ |
| 4 | $\Delta r = -0.0028$, 95% CI [-0.0064, 0.0010], $d = -0.2451$, $p = .9244$ | $\Delta r = -0.0006$, 95% CI [-0.0049, 0.0042], $d = -0.0594$, $p = .5891$ | $\Delta r = -0.0002$, 95% CI [-0.0030, 0.0027], $d = -0.0364$, $p = 0.5616$ | $\Delta r = -0.0026$, 95% CI [-0.0056, 0.0006], $d = -0.3914$, $p = 0.9319$ |
| 5 | $\Delta r = -0.0015$, 95% CI [-0.0040, 0.0011], $d = -0.1986$, $p = .8802$ | $\Delta r = -0.0019$, 95% CI [-0.0052, 0.0011], $d = -0.2896$, $p = .8723$ | $\Delta r = -0.0028$, 95% CI [-0.0055, -0.0002], $d = -0.4819$, $p = 0.9719$ | $\Delta r = -0.0025$, 95% CI [-0.0051, 0.0003], $d = -0.4265$, $p = 0.9535$ |
| 7 | $\Delta r = 0.0007$, 95% CI [-0.0025, 0.0041], $d = 0.0651$, $p = .3592$ | $\Delta r = 0.0010$, 95% CI [-0.0036, 0.0060], $d = 0.0968$, $p = .3524$ | $\Delta r = -0.0012$, 95% CI [-0.0042, 0.0023], $d = -0.1679$, $p = 0.7551$ | $\Delta r = -0.0024$, 95% CI [-0.0052, 0.0004], $d = -0.3821$, $p = 0.9300$ |
| 8 | $\Delta r = -0.0004$, 95% CI [-0.0031, 0.0021], $d = -0.0558$, $p = .6333$ | $\Delta r = 0.0011$, 95% CI [-0.0015, 0.0039], $d = 0.1914$, $p = .2289$ | $\Delta r = 0.0044$, 95% CI [0.0006, 0.0082], $d = 0.5364$, $p = 0.0210$ | $\Delta r = 0.0009$, 95% CI [-0.0012, 0.0032], $d = 0.1884$, $p = 0.2215$ |

**S4 Table. Peak coordinates and Glasser atlas labels for clusters surviving voxelwise thresholding ($p < .01$, one-tailed; uncorrected, $k \geq 20$ voxels) in the whole-brain searchlight analysis of hexadirectional modulation.**

| Region | Volume (cubic mm) | MNI x | MNI y | MNI z | Peak $t$ | Glasser parcel | % cluster in parcel |
|---|---|---|---|---|---|---|---|
| Cblm_CrusII_L | 1608 | -29 | -69 | -43 | 7.03 | Cerebellum | 68 |
| Ctx_STSvp_L | 1592 | -55 | -33 | -9 | 6.38 | Cortex_Default_ModeB | 29 |
| Ctx_p32pr_L | 2024 | -19 | 14 | 36 | 5.98 | Cortex_Ventral_AttentionA | 1 |
| Ctx_TE2a_R | 1152 | 52 | -5 | -33 | 5.54 | Cortex_Limbic | 57 |
| Ctx_3b_L | 3424 | -21 | -29 | 56 | 5.35 | Cortex_SomatomotorA | 14 |
| Cau_L | 1480 | -27 | -11 | 26 | 5.30 | Basal_ganglia | 23 |
| Cblm_V_R | 776 | 28 | -43 | -25 | 5.24 | Cerebellum | 72 |
| Cblm_VI_R | 616 | 10 | -67 | -25 | 5.03 | Cerebellum | 61 |

| | | | | | | | |
|---|---|---|---|---|---|---|---|
| Ctx_4_R | 2584 | 44 | -17 | 30 | 4.90 | Cortex_SomatomotorA | 11 |
| GPi_L | 920 | -15 | -9 | -7 | 4.85 | Basal_ganglia | 44 |
| Ctx_11l_R | 1696 | 20 | 42 | -15 | 4.83 | Cortex_Fronto_ParietalB | 27 |
| Cblm_VIIIa_L | 2592 | -23 | -63 | -59 | 4.81 | Cerebellum | 65 |
| Cblm_I_IV_R | 816 | 6 | -45 | -17 | 4.80 | Cerebellum | 80 |
| Ctx_6r_R | 2768 | 42 | 8 | 28 | 4.80 | Cortex_Ventral_AttentionA | 17 |
| Ctx_4_R | 3696 | 14 | -25 | 66 | 4.79 | Cortex_SomatomotorA | 20 |
| Ctx_V1_R | 640 | 20 | -77 | 14 | 4.79 | Cortex_Visual_Peripheral | 46 |
| Ctx_pOFC_L | 1376 | -17 | 16 | -17 | 4.78 | Cortex_Limbic | 31 |
| Ctx_TGv_R | 1680 | 38 | 6 | -47 | 4.74 | Cortex_Limbic | 49 |
| Ctx_d32_R | 2160 | 14 | 40 | 16 | 4.67 | Cortex_Ventral_AttentionB | 21 |
| Ctx_6mp_L | 560 | 2 | -15 | 60 | 4.67 | Cortex_SomatomotorA | 34 |
| Ctx_POS2_R | 656 | 14 | -65 | 28 | 4.63 | Cortex_Fronto_ParietalC | 67 |
| Ctx_PCV_R | 640 | 16 | -51 | 46 | 4.59 | Cortex_Fronto_ParietalC | 16 |
| Ctx_V3_L | 4160 | -23 | -87 | 18 | 4.58 | Cortex_Visual_Central | 17 |
| Ctx_2_L | 1368 | -27 | -47 | 60 | 4.58 | Cortex_Dorsal_AttentionB | 25 |
| Bstem_Midbrv_R | 576 | 10 | -7 | -17 | 4.54 | Brainstem | 29 |
| Ctx_V4_L | 880 | -29 | -67 | -5 | 4.50 | Cortex_Visual_Central | 43 |
| Ctx_10v_R | 2464 | -1 | 44 | -21 | 4.44 | Cortex_Default_ModeA | 22 |
| Ctx_10v_R | 1416 | 10 | 62 | -3 | 4.43 | Cortex_Default_ModeA | 31 |
| Ctx_V3_R | 2040 | 22 | -91 | 12 | 4.38 | Cortex_Visual_Central | 18 |
| Ctx_PF_R | 1072 | 64 | -45 | 34 | 4.38 | Cortex_Ventral_AttentionA | 48 |
| Ctx_V4_R | 1200 | 36 | -83 | -11 | 4.36 | Cortex_Visual_Central | 44 |
| Bstem_Ponscd | 816 | 14 | -37 | -35 | 4.34 | Brainstem | 95 |
| Putamen_Pp_L | 808 | -25 | -1 | -13 | 4.31 | Basal_ganglia | 32 |
| Ctx_46_R | 936 | 30 | 34 | 36 | 4.31 | Cortex_Ventral_AttentionB | 23 |
| Ctx_OP1_L | 1144 | -51 | -23 | 28 | 4.27 | Cortex_SomatomotorB | 10 |

| | | | | | | | |
|---|---|---|---|---|---|---|---|
| Cblm_VIIb_R | 568 | 34 | -65 | -49 | 4.27 | Cerebellum | 75 |
| Ctx_1_L | 1376 | -57 | -21 | 52 | 4.26 | Cortex_SomatomotorA | 66 |
| Ctx_PeEc_L | 600 | -31 | -15 | -37 | 4.09 | Cortex_Limbic | 51 |
| Ctx_8BL_R | 656 | 12 | 44 | 54 | 4.02 | Cortex_Default_ModeB | 76 |
| Ctx_6d_L | 808 | -27 | -17 | 74 | 4.00 | Cortex_SomatomotorA | 49 |
| Ctx_DVT_R | 1200 | 28 | -67 | 18 | 3.99 | Cortex_Fronto_ParietalC | 20 |
| No label | 592 | -29 | 44 | -1 | 3.97 | No description | 0 |
| Ctx_a24_L | 808 | -5 | 38 | 2 | 3.96 | Cortex_Default_ModeA | 22 |
| Ctx_V1_L | 2120 | -13 | -79 | 6 | 3.93 | Cortex_Visual_Peripheral | 29 |
| Ctx_STV_R | 632 | 52 | -41 | 24 | 3.92 | Cortex_Temporal_Parietal | 18 |
| No label | 600 | 20 | 16 | 40 | 3.92 | No description | 0 |
| Cblm_VI_R | 1384 | 8 | -75 | -11 | 3.92 | Cerebellum | 41 |
| Caudate_Ca_R | 720 | 12 | 14 | 10 | 3.86 | Basal_ganglia | 74 |
| Ctx_9_46d_R | 1040 | 32 | 44 | 16 | 3.81 | Cortex_Ventral_AttentionB | 6 |
| GPe_R | 1360 | 20 | 2 | -3 | 3.76 | Basal_ganglia | 38 |
| Ctx_RI_R | 704 | 34 | -25 | 22 | 3.75 | Cortex_SomatomotorB | 31 |
| Ctx_OP4_L | 592 | -55 | -7 | 12 | 3.66 | Cortex_SomatomotorB | 45 |
| Ctx_SCEF_R | 792 | 8 | -5 | 62 | 3.65 | Cortex_Ventral_AttentionA | 68 |
| Ctx_1_L | 1504 | -45 | -21 | 64 | 3.62 | Cortex_SomatomotorA | 32 |
| Ctx_V1_L | 640 | -13 | -91 | 10 | 3.41 | Cortex_Visual_Peripheral | 32 |
| Ctx_7Am_L | 576 | -1 | -51 | 62 | 3.32 | Cortex_Dorsal_AttentionB | 74 |
| Ctx_PH_R | 632 | 54 | -71 | -11 | 3.12 | Cortex_Dorsal_AttentionA | 44 |
